# Structural organisation of perisomatic inhibition in the mouse medial prefrontal cortex

**DOI:** 10.1101/2023.06.14.545032

**Authors:** Petra Nagy-Pál, Judit M. Veres, Zsuzsanna Fekete, Mária R. Karlócai, Filippo Weisz, Bence Barabás, Zsófia Reéb, Norbert Hájos

**Affiliations:** ELRN Institute of Experimental Medicine, 1083, Budapest, Hungary; János Szentágothai School of Neurosciences, Semmelweis University, 1085, Budapest, Hungary; Doctoral School of Biology, Institute of Biology, ELTE Eötvös Loránd University, 1117 Budapest, Hungary; The Linda and Jack Gill Center for Molecular Bioscience, Indiana University Bloomington, 47405, Indiana, USA; Program in Neuroscience, Department of Psychological and Brain Sciences, Indiana University Bloomington, 47405, Indiana, USA

## Abstract

Perisomatic inhibition profoundly controls neural function. However, the structural organisation of inhibitory circuits giving rise to the perisomatic inhibition in the higher-order cortices is not completely known. Here, we performed a comprehensive analysis of those GABAergic cells in the medial prefrontal cortex (mPFC) that provide inputs onto the somata and proximal dendrites of pyramidal neurons. Our results show that most GABAergic axonal varicosities contacting the perisomatic region of superficial (layer 2/3) and deep (layer 5) pyramidal cells express parvalbumin (PV) or cannabinoid receptor type 1 (CB1). Further, we found that the ratio of PV/CB1 GABAergic inputs is larger on the somatic membrane surface of pyramidal tract neurons in comparison to those projecting to the contralateral hemisphere. Our morphological analysis of in vitro labelled PV+ basket cells (PVBC) and CCK/CB1+ basket cells (CCKBC) revealed differences in many features. PVBC dendrites and axons arborized preferentially within the layer where their soma was located. In contrast, the axons of CCKBCs expanded throughout layers, though their dendrites were found preferentially either in superficial or deep layers. Finally, using anterograde trans-synaptic tracing we observed that PVBCs are preferentially innervated by thalamic and basal amygdala afferents in layer 5a and 5b, respectively. Thus, our results suggest that PVBCs can control the local circuit operation in a layer-specific manner via their characteristic arborization, while CCKBCs rather provide cross-layer inhibition in the mPFC.

**Significance Statement:** Inhibitory cells in cortical circuits are crucial for the precise control of local network activity. Nevertheless, in higher-order cortical areas that are involved in cognitive functions like decision making, working memory and cognitive flexibility, the structural organisation of inhibitory cell circuits is not completely understood. In this study we show that perisomatic inhibitory control of excitatory cells in the medial prefrontal cortex is performed by two types of basket cells endowed with different morphological properties that provide inhibitory inputs with distinct layer specificity on cells projecting to disparate areas. Revealing this difference in innervation strategy of the two basket cell types is a key step towards understanding how they fulfil their distinct roles in cortical network operations.

## Introduction

The medial part of the PFC (mPFC) has a critical role in controlling various higher order cognitive functions (Goldman-Rakic, 1988; Miller, 2000; Fuster, 2006), such as working memory, attention, decision making or fear behaviour (Cohen et al., 1996; Euston et al., 2012; Tovote et al., 2015; Lee and Seo, 2016; Uddin, 2021), yet our knowledge about the circuit organisation within this cortical area is still limited. The mPFC networks are composed of excitatory glutamatergic pyramidal neurons and a variety of GABAergic inhibitory cells (DeFelipe and Farinas, 1992; Anastasiades and Carter, 2021) . Although the inhibitory neurons constitute only 10-25% of the whole neuronal population in the mPFC (Markram et al., 2004; Xu et al., 2019; Le Merre et al., 2021), they are able to effectively modulate and control the activity of large pyramidal cell populations (Isaacson and Scanziani, 2011; Kepecs and Fishell, 2014; Pelkey et al., 2017), to synchronize pyramidal cell firing at different frequencies (Buzsaki and Chrobak, 1995; Jung and Carlen, 2021), to channel the information flow within the local circuits (Reyes et al., 1998; Pouille and Scanziani, 2004; Isaacson and Scanziani, 2011), and to promote the storage of information (Frankland and Bontempi, 2005; Topolnik and Tamboli, 2022).

Previous studies have shown that, similarly to other cortical regions, several different GABAergic cell types are present in the mPFC (Kawaguchi and Kubota, 1997; Markram et al., 2004; Jiang et al., 2015; Tremblay et al., 2016). Different inhibitory interneurons innervate distinct, often non-overlapping membrane domains of postsynaptic neurons, allowing them to fulfil specific roles in microcircuit function (Miles et al., 1996; Larkum et al., 1999; Petilla Interneuron Nomenclature et al., 2008; Larkum, 2013; Kepecs and Fishell, 2014). Perisomatic region targeting interneurons innervate mostly the somata, the proximal dendrites and axon initial segment of pyramidal cells. As the synaptic inhibition of these membrane surfaces controls the spike generation very efficiently (Cobb et al., 1995; Miles et al., 1996; Veres et al., 2014; Veres et al., 2017), inhibitory cells innervating the perisomatic region are well suited to regulate the discharge of neurons (Cobb et al., 1995; Miles et al., 1996; Somogyi et al., 1998; Kawaguchi and Kondo, 2002; Freund and Katona, 2007). In cortical structures two types of basket cells, expressing either PV or CCK/CB1, give rise to innervation of the somata and proximal dendrites of pyramidal cells (Somogyi et al., 1998; Freund and Katona, 2007; Kepecs and Fishell, 2014). Although these basket cells target the same membrane surface, they are distinct in their morphological features and electrophysiological characteristics (Hefft and Jonas, 2005; Klausberger et al., 2005; Glickfeld and Scanziani, 2006; Foldy et al., 2007; Szabo et al., 2010; Vereczki et al., 2016; Barsy et al., 2017) as it has been found in other cortical structures suggesting that they play distinct roles in network operation (Freund, 2003). At present, however, our knowledge is limited regarding the function of CCKBCs in comparison to PVBCs in the mPFC. By revealing how basket cell networks are structurally organised in the mPFC, we can predict their function at the microcircuit level.

In this study, we first defined the layers within the mouse mPFC which helped us to investigate the organisation of perisomatic inhibition in a layer-dependent manner. Next, we determined the source and the ratio of GABAergic inputs innervating the perisomatic region of pyramidal cells. Then, we characterized and compared the morphological features of interneurons providing the vast majority of perisomatic inputs on pyramidal cells. We observed morphologically distinct subgroups of basket cells based on the differences in their input-output properties. Finally, we examined whether the extra-prefrontal cortical afferents target PV-containing interneurons in the different layers with distinct likelihood. Based on the morphological differences, PVBCs in the mPFC can accomplish layer-specific regulation of pyramidal cell operation, while CCKBCs may control pyramidal cell function more broadly.

## Materials and methods

### Experimental Design

#### Experimental animals

Experiments were approved by the Committee of the Scientific Ethics of Animal Research (22.1/360/3/2011) and all procedures involving animals were performed according to methods approved by Hungarian legislation (1998 XXVIII. section 243/1998, renewed in 40/2013.) and institutional guidelines of ethical code. All procedures complied with the European convention for the protection of vertebrate animals used for experimental and other scientific purposes (Directive 86/609/CEE and modified according to the Directives 2010/63/EU). Every effort was taken to minimise animal suffering and the number of animals used. Adult mice (p50-140) were used from the following lines: C57BL/6J, FVB/Ant-Fx (wild types, Charles River), BAC-PV-eGFP (Meyer et al., 2002), BAC-CCK-DsRed (Mate et al., 2013), Ai6 reporter line (CAG-LSL-ZsGreen1, #007906, www.jax.org) and Pvalb-IRES-Cre (PV-Cre, #017320, www.jax.org).

### Surgical procedures

#### Retrograde cell labelling

Anaesthesia was induced with 125 mg/kg ketamine and 5 mg/kg xylazine. Wild type mice from both sexes (n=2-2 for basal amygdala (BA) and periaqueductal grey (PAG) injection and n=5-5 for contralateral PFC (cPFC) and dorsal striatum (DS) injection) were secured in a stereotaxic frame and unilateral or bilateral injections of fluorogold or choleratoxin B subunit tracers dissolved in glycerol (FG 2% iontophoresis by 2 µA pulses with 2/2 s on/off duty cycle for 5 minutes and CTB 0.5% iontophoresis by 5 µA pulses with 2/2 s on/off duty cycle for 7-10 minutes) or retrograde pAAVrg-CAG-GFP (Addgene: 37825-AAVrg) or retrograde AAVrg-EF1a-mCherry-IRES-Flpo (Addgene: 55634-AAVrg) viruses (200 nl with 3nl/sec flow rate) were aimed at the following coordinates from Bregma (in cm): for BA injections: AP -0.15, ML 0.31, DV 0.43; for (DS) injections: AP 0.06-0.07, ML 0.13, DV 0.23; for PAG injections: AP -0.46-0.49, ML 0.05, DV 0.11-0.12; for cPFC injections: AP 0.15-0.18, ML next to the sinus, DV 0.1-0.15. After 4-8 days of tracer injection and after 4 weeks of virus injection, mice were transcardially perfused with 4% paraformaldehyde (PFA) in 0.1M phosphate buffer (PB) for 30-40 min and PFC containing sections of 50-100 μm thickness were prepared using a Vibratome (Leica) and stored in 0.1M PB with 0.05% Na-azide until further processing.

### Anterograde trans-synaptic viral labelling

Anaesthesia was induced with 125 mg/kg ketamine and 5 mg/kg xylazine. Male PV-Cre and both male and female Ai6 mice were secured in a stereotaxic frame and bilateral injections of 200-300 nl AAV1 virus vectors (AAV1-EF1a-DIO-ChETA-eYFP (Addgene: 26968) to PV-Cre mice and pENN-AAV1-hSyn-Cre-WPRE-hGH (Addgene: 105553) to Ai6 mice) were aimed with 3nl/sec flow rate at the following coordinates from Bregma (in cm): for BA injections AP -0.15, ML 0.32-0.35, DV 0.44-0.50; for midline thalamus injections AP -0.12-0.13, ML next to the sinus, DV 0.30-0.33; for lateral entorhinal cortex (LEnt) injections AP - 0.42, ML 0.375, DV 0.28. After 4 weeks of recovery, mice were transcardially perfused with 4% PFA in 0.1M PB for 30-40 minutes and PFC sections were prepared as described above.

### Tissue processing and immunocytochemistry

All anatomical data, including those acquired with viral vectors and retrograde tracers, were obtained from immunostained brain slices. After perfusion 50-100 μm thick sections were prepared using a Vibratome (Leica VT100S). Slices were thoroughly washed in 0.1M PB for several times (4-5 times for 10-15 minutes). For fluorescent labelling, slices were blocked with 10% Normal Donkey Serum (NDS, Vector Laboratories) or 10% Normal Goat Serum (NGS, Vector Laboratories) and 0.5% TritonX-100 in 0.1M PB for 30-60 minutes at room temperature. Then, sections were incubated in a mixture of different primary antibodies diluted in PB containing 2% NDS or 2% NGS, 0.5% or 2% TritonX-100 and 0.05% Na-azide overnight at room temperature for an additional 3-6 days at 4°C. The applied primary antibodies are detailed in Table 1 grouped and numbered by the experiments: for the determination of the borders of PFC layers (1); for the visualisation of GABAergic inputs on the perisomatic region (2); for the visualisation of inputs onto retrogradely labelled cells (3-6); for validating the CCK- or PV-content in the transgenic mice (7-8); for the examination of CB1 receptor content of visualised CCKBCs (9); for the target distribution of CCKBCs and PVBCs on random PCs (10); for the visualisation of the monosynaptic input-receiving PV-containing cells (11-12). After washing out primary antibodies several times, slices were treated with secondary antibodies diluted in 0.1M PB and 1% NDS or NGS for 2-4 hours (Table 1). Following several washes in PB, sections were mounted on glass slides in Vectashield (Vector Laboratories).

**Table 1.**
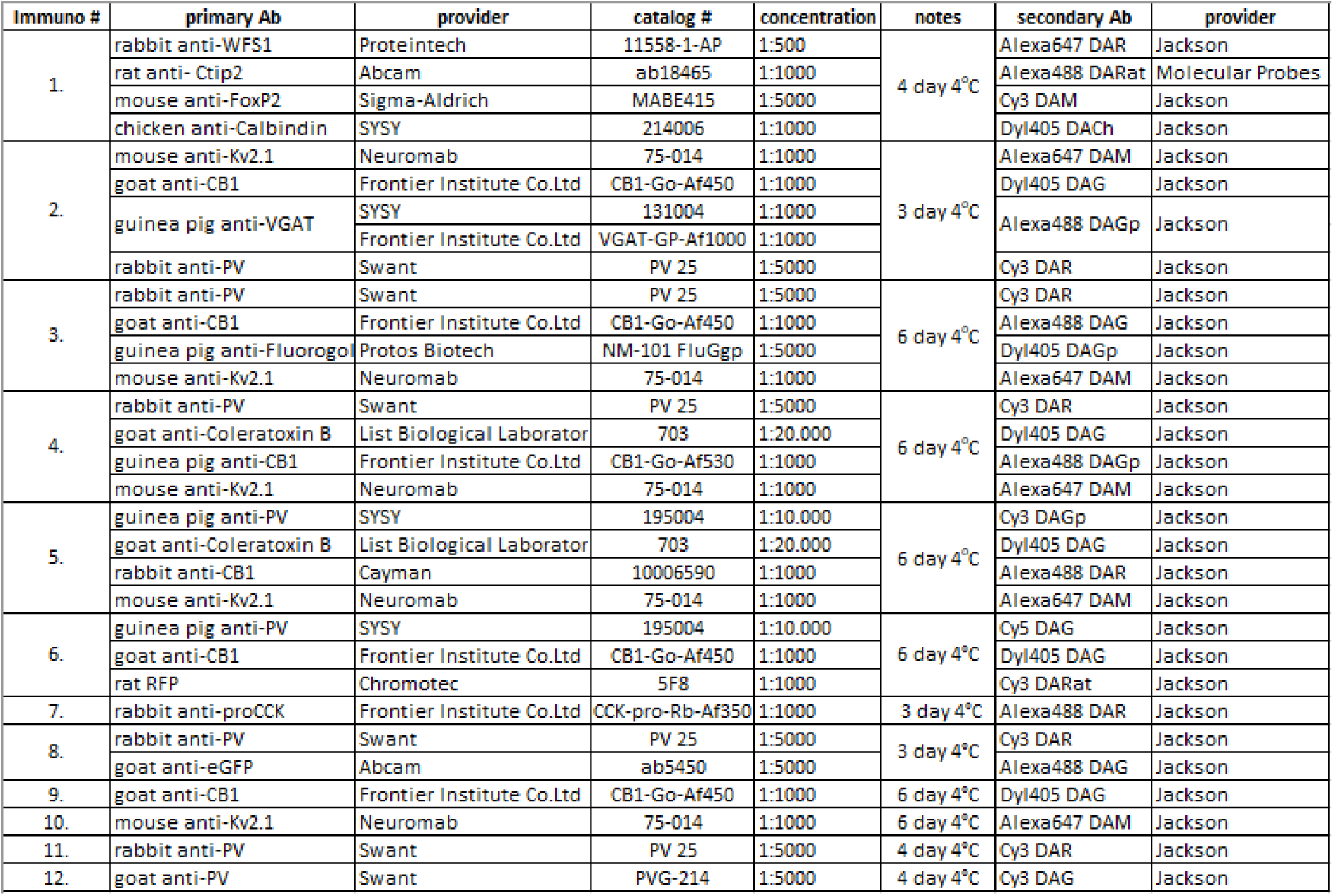
Antibodies used in anatomical experiments. Abbreviation of secondary antibodies: DAR: donkey anti rabbit, DARat: donkey anti rat, DAM: donkey anti mouse, DACh: donkey anti chicken, DAG: donkey anti goat, DAGp: donkey anti guinea pig. All the secondary antibodies were used in 1:500 dilution.

### Interneuron recording, labelling *in vitro*

*In vitro* biocytin labelling of interneurons was carried out as described earlier in detail (Veres et al., 2014). Briefly, CCK-DsRed and PV-eGFP mice were deeply anesthetized with isoflurane and decapitated. The brain was quickly removed and placed into ice-cold cutting solution containing (in mM): 252 sucrose, 2.5 KCl, 26 NaHCO_3_, 1 CaCl_2_, 5 MgCl_2_, 1.25 NaH_2_PO_4_, 10 glucose, bubbled with 95% O_2_ and 5% CO_2_ (carbogen gas). Coronal slices of 200-μm thickness containing the PFC region were prepared with a Leica VT1200S Vibratome (Wetzlar, Germany), and kept in an interface-type holding chamber containing artificial cerebrospinal fluid (ACSF) at 36 °C that gradually cooled down to room temperature. ACSF contained (in mM) 126 NaCl, 2.5 KCl, 1.25 NaH_2_PO_4_, 2MgCl_2_, 2 CaCl_2_, 26 NaHCO_3_, and 10 glucose, bubbled with carbogen gas. Interneurons were selected based on the presence of the fluorescent proteins (DsRed or eGFP) excited by an UV lamp and visualised by a CCD camera (Hamamatsu Photonics or Andor Zyla). Targeted cells were recorded under visual guidance using differential interference contrast microscopy (Olympus BX61W or Nikon FN-1) that laid 50–100 μm below the surface of the acute slice. Interneurons were recorded in whole-cell mode using a K-gluconate based intrapipette solution containing biocytin to label their processes [intrapipette solution (in mM): 110 K-gluconate, 4 NaCl, 2 Mg-ATP, 20 HEPES, 0.1 EGTA, 0.3 GTP (sodium salt), 10 phosphocreatine, and 0.2% biocytin adjusted to pH 7.3 using KOH and with an osmolarity of 290 mOsm/L]. After the recordings, slices were fixed in 4% PFA and Alexa488-coupled streptavidin (1:10,000 in TBS, Molecular Probes) or Alexa647-coupled streptavidin (1:10,000 in TBS, Molecular Probes) was used to visualise the fine details of the neurons in the entire slice.

### Images and analysis

Fluorescent images were taken with a Nikon A1R or C2 confocal laser scanning microscope (Nikon Europe, Amsterdam, The Netherlands) using distinct settings for different objectives: CFI Super Plan Fluor 20X, N.A. 0.45, z step size: 1 μm, xy: 0.62 μm/pixel for the cell reconstruction; CFI Plan Apo VC10X, N.A. 0.30, single plane, xy: 0.31 μm/pixel for the determination of the borders of layers within the mPFC and for the validation of reporter protein content of animals; CFI Plan Apo VC60X Oil objective, N.A. 1.40, z step size: 0.13 μm, xy: 0.08 μm/pixel for the visualisation of GABAergic inputs on the perisomatic region and for the analysis of the density of CB1 and PV boutons on the retrogradely labelled cells; CFI Plan Apo VC60X Oil objective, N.A. 1.40, z step size: 0.2, xy: 0.21 μm/pixel for the analysis of target distribution of basket cells on random pyramidal cells; CFI Plan Apo VC10X, N.A. 0.30, z step size: 3 μm, xy: 0.63 μm/pixel for the visualisation of the monosynaptic input receiving PV-containing cells.

Reconstruction and analysis of the 3D confocal images was performed with the Neurolucida 10.53 software (MBF Bioscience) and NIS Elements software (Nikon). The properties of axonal and dendritic arbours and surface analysis were performed with Neurolucida Explorer software (MBF Bioscience). Values were corrected for shrinkage and flattening of the tissue (x, y and z axis correction on pictures taken by using CFI Plan Apo VC60X Oil objective: 1.08; x, y axis correction was 1 and z axis correction was 2.5 on pictures taken from the biocytin-labelled basket cells by using CFI Super Plan Fluor 20X objective). Schematic representation of brain slices from thalamus injection and PFC were achieved using Inkscape (Free Software Foundation Inc.) open access program.

### Quantification of inputs on Kv2.1-immunolabelled somata

During quantification, different aspects were taken into account. Boutons were considered putative contacts if no apparent gap was visible between the labelled bouton and the surface of Kv2.1-stained somata when examined in 3D view of confocal images. Varicosities located at branch points were not counted as putative contacts, as it has been shown by electron microscopy that these boutons do not form synapses (Veres et al., 2014; Veres et al., 2017).

### Statistical and Cluster Analysis

For comparison of data with normal distribution according to the Shapiro-Wilk test, the two sample T-test and ANOVA were used. For data with non-normal distribution the Mann-Whitney U test (M-W test), Wilcoxon Signed Rank test and Kruskal-Wallis ANOVA (K-W ANOVA) was used. For *post hoc* analysis Dunn’s test or M-W test was used. For the comparison of distributions, the two sample Kolmogorov-Smirnov test (K-S test) and Chi-Square Homogenity test were used. All statistics were performed using Origin 8.6 or 9.2 (Northampton, MA) or online LibreTexts statistical programs (www.stats.libretexts.org). Exact p values were indicated when p was higher than 0.001 considering the rounding rules. Data are presented as mean ± SEM, unless indicated otherwise. The cluster and principal component analysis (PCA) were performed using Origin 8.6 or 9.2. The PCA showed the main components from the 5 axonal and 5 dendritic distribution ratios between layers, and cluster analysis applying Ward methods was made based on these selected main components.

## Results

### Defining the layers in the mPFC

As the layers in the mPFC are not labelled consistently across studies (Clarkson et al., 2017; Lu et al., 2017; Anastasiades et al., 2018), we first defined the distinct layers by visualising a combination of markers that have already proven to be good tools for differentiating layers in other cortical areas (Figure 1A) (Cruikshank et al., 2001; Arlotta et al., 2005; Luuk et al., 2008; Hisaoka et al., 2010). Our study was conducted primarily in the prelimbic area of the mouse mPFC, but, as the borders between this area and the neighbouring cortical regions (the anterior cingulate cortex and the infralimbic cortex) are ill-defined, we will refer to the investigated area as the mPFC. To visualise the borders between the two superficial layers, layers 2 and 3, the transcription factor wolfram syndrome 1 protein (WFS1) and the calcium-binding protein Calbindin (Calb) were used. WFS1 revealed the neurons in layer 2 (Luuk et al., 2008) (Figure 1B_1_-B_2_), while Calb was expressed at high levels in both layers 2 and 3 (Cruikshank et al., 2001) (Figure 1C_1_-C_2_). We surprisingly observed that in the dorsal part of the mPFC, these two layers fuse and form a narrow layer 2/3 (Figure 1B_2_-C_2_), preventing the separation of these two superficial layers. To determine the borders of deep layers, two additional transcription factors were applied. COUP-TF interacting protein 2 (Ctip2) was found to be present in neurons located in layer 5b and layer 6 (Arlotta et al., 2005) (Figure 1D_1_-D_2_), while Forkhead box protein P2 (FoxP2) was expressed predominantly in layer 6 neurons (Hisaoka et al., 2010) (Figure 1E_1_-E_2_). As the layer 4 cannot be defined in the rodent PFC (Uylings et al., 2003), the borders between layers in the mPFC is clearly determined by the visualisation of these markers (Figure 1F), allowing us to investigate the structural organisation of perisomatic inhibition in a layer-dependent manner

**Figure 1:**
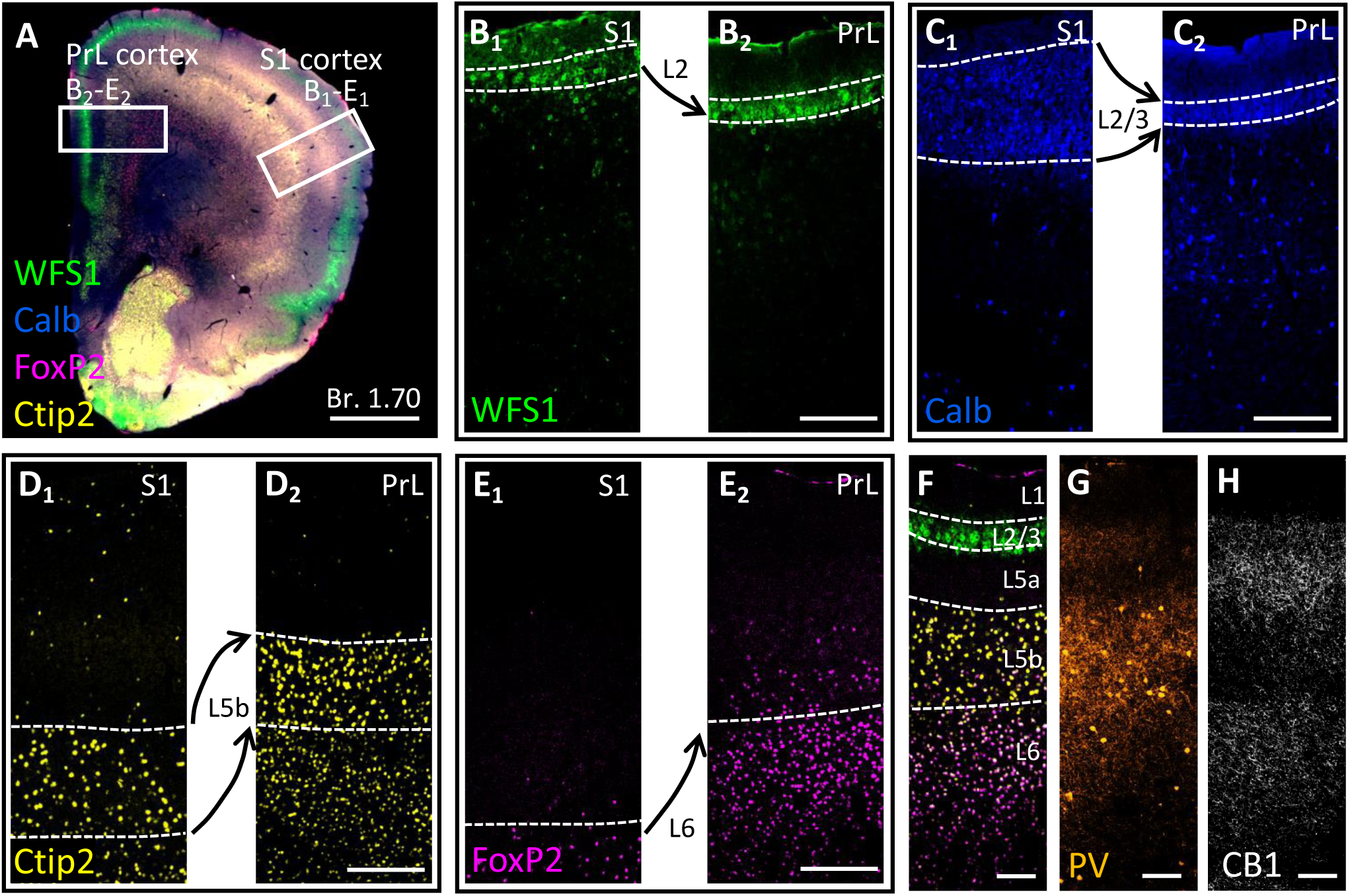
Defining the layers in the mPFC by using antibodies developed against the Ca^2+^ binding protein calbindin (Calb) and transcription factors (WFS1, Ctip2, FoxP2) (A) A low magnification multicolour confocal image taken from a mPFC section immunostained against WFS1, Calb, Ctip2 and FoxP2. White empty rectangles indicate the prelimbic (PrL) subregion of the mPFC and the primary somatosensory cortex (S1), which are shown at a higher magnification in B-E. Scale bar: 1 mm. (B-E) Higher magnification confocal images taken from the S1 and PrL region, and immunolabelled for WFS1, Calb, Ctip2, FoxP2. Dashed white lines indicate the boundaries of the layers defined by the immunolabeling. (F) A multicolour confocal image taken from a PrL region that was immunostained against WFS1 (green), Ctip2 (yellow) and FoxP2 (magenta), indicating the clear boundaries of the layers. (G, H) Immunostaining against PV and CB1 in the PrL region of mPFC. Scale bar: 100 µm.

### The vast majority of perisomatic GABAergic inputs originate from CB1- and PV-expressing axonal varicosities

The perisomatic membrane surface of neurons is a distinctive site, where the action potential generation can be controlled powerfully by inhibitory GABAergic synapses (Cobb et al., 1995; Miles et al., 1996; Veres et al., 2017). As prior studies elucidated, mostly PVBCs and CCKBCs provide these inhibitory axon terminals onto the perisomatic region of pyramidal cells in cortical areas studied so far (Freund and Katona, 2007; Takacs et al., 2015; Vereczki et al., 2016). However, at present it is unclear how the perisomatic inhibition originating from basket cells expressing PV or CB1 is organized in the mPFC at the structural level. Immunostaining against PV and CB1 revealed that the distribution of axonal boutons expressing these proteins varied among the layers in the mPFC (Figure 1G-H), implying that pyramidal cell in different layers may receive distinct ratio of perisomatic inputs from PVBCs and CCKBCs. To support this hypothesis, we first assessed the sources of perisomatic GABAergic inputs in the distinct layers of mPFC using multichannel high-resolution confocal microscopic investigations. The perisomatic region of neurons was visualised by immunostaining against the voltage-gated potassium channel subunit Kv2.1 (Vereczki et al., 2016), while the vesicular GABA transporter (VGAT) was used to reveal GABAergic axonal varicosities and their neurochemical content by immunostaining against PV and CB1 (Figure 2A). Our analysis uncovered that the vast majority of the VGAT-immunolabelled boutons were immunopositive for PV or CB1 in layer 2/3, 5a and 5b (Figure 2B). Interestingly, we found that neurons in layer 5b receive a significantly larger PV+/CB1+ perisomatic input ratio than neurons in layer 2/3 and layer 5a (Figure 2D-E). As pyramidal tract (PT) neurons projecting to subcortical targets are located predominantly in layer 5b in the mPFC (Gabbott et al., 2005; Kawaguchi, 2017; Anastasiades and Carter, 2021), our observations may indicate that depending on the target area of pyramidal cell axons, they could be innervated by various ratios of PV+/CB1+ boutons (Bodor et al., 2005; Varga et al., 2010) in a layer-selective manner (Figure 1G, H). To directly test this hypothesis, we examined the CB1+ and PV+ axonal varicosities on the Kv2.1-labelled perisomatic region of pyramidal cells projecting to the basal amygdala (BA), contralateral PFC (cPFC), dorsal striatum (DS) or periaqueductal grey (PAG) (Figure 3). The soma location of pyramidal cells projecting to these remote areas were similar to that described earlier (Gabbott et al., 2005; Lu et al., 2017; Anastasiades and Carter, 2021) (Figure 3A-B). The analysis of the ratio of PV+/CB1+ boutons forming close appositions on the perisomatic membrane surface of pyramidal cells projecting to distinct regions revealed that the proportion of these two GABAergic inputs was similar in layer 2/3 and 5a regardless of where pyramidal neurons projected (Figure 3F). This ratio, however, showed a difference in layer 5b and was higher on PAG-projecting pyramidal cells compared to cPFC-projecting pyramidal cells (Figure 3F).

**Figure 2.**
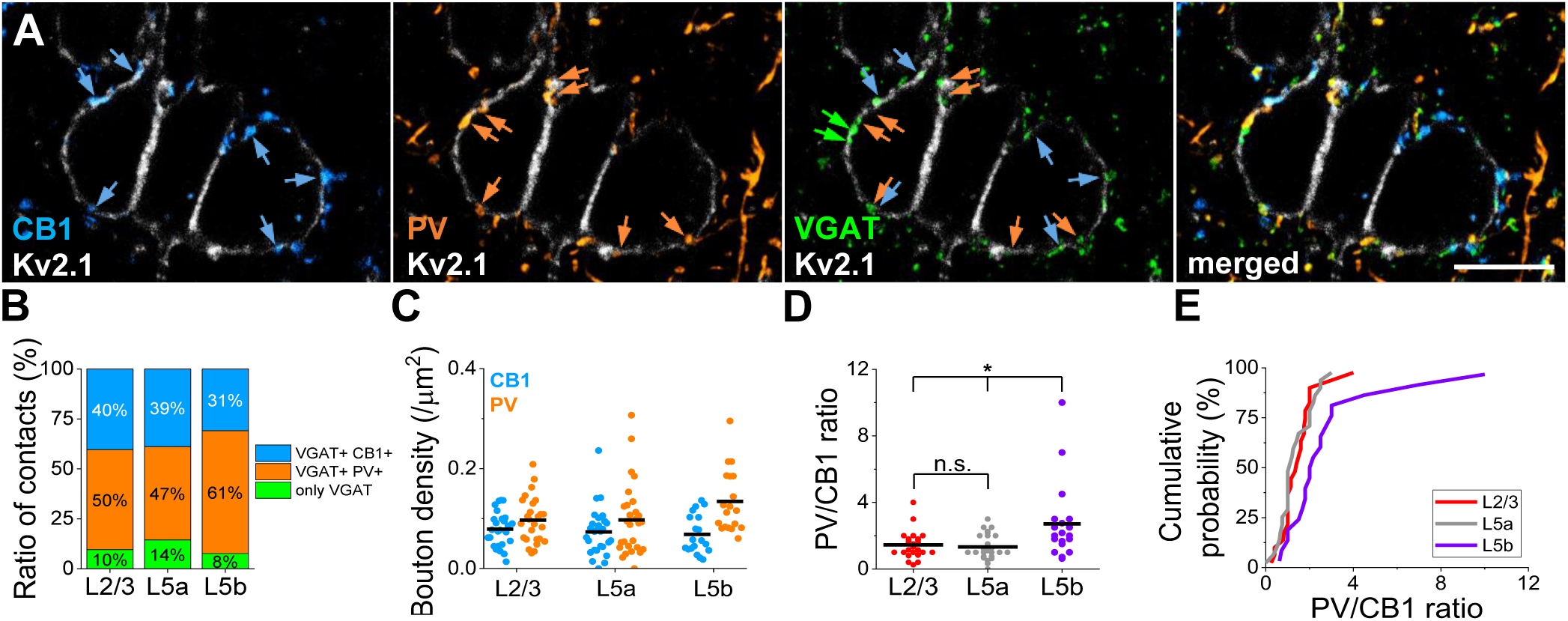
The vast majority of perisomatic GABAergic inputs originate from CB1- and PV-expressing boutons. (A) Multicolour maximum z-intensity projection confocal images taken of sections immunostained for Kv2.1, CB1, PV and VGAT. Blue arrows indicate the CB1 and VGAT co-expressing boutons on Kv2.1-labelled profiles, orange arrows point to PV and VGAT-co-expressing terminals that are in close appositions with Kv2.1-immunostained soma membranes, while green arrows show VGAT-immunopositive boutons that lack PV or CB1 immunoreactivity on Kv2.1-labelled somata. Scale bar: 10 µm. (B) The ratio of boutons expressing CB1, PV and VGAT only on the Kv2.1-immunostained somata in three different layers of the mPFC. (L2/3: only VGAT: 9.61 ± 1.88%, VGAT+ PV+: 50.02 ± 2.79%, VGAT+ CB1+: 40.37 ± 2.44%; L5a: only VGAT: 14.47 ± 2.6%, VGAT+ PV+: 46.69 ± 3.5%, VGAT+ CB1+: 38.84 ± 2.94%; L5b: VGAT: 7.69 ± 1.64%, VGAT+ PV+: 61.43 ± 3.46%, VGAT+ CB1+: 30.88 ± 2.92%) (C) CB1 and PV bouton density on the surface of Kv2.1-labelled somata in different layers. Black lines represent the mean in each case. (Mean (bouton/µm^2^): L2/3 CB1: 0.0787; L2/3 PV: 0.0968; L5a CB1: 0.0731; L5a PV: 0.0971; L5b CB1: 0.0683; L5b PV: 0.1341.) (D) The ratio of PV and CB1-expressing boutons on Kv2.1-labelled membrane profiles in distinct layers. Each dot represents the ratio which was determined on single pyramidal neurons, while black lines show the mean. In layer 2/3 n=26, in layer 5a n=28, while in layer 5b n=19 somata were examined. (One way ANOVA: p=0.0022; Two sample t test: L2/3 vs. L5a: p=0.59; L2/3 vs. L5b: p=0.012; L5a vs. L5b: p=0.0061. Mean: L2/3: 1.442; L5a: 1.327; L5b: 2.712.) (E) Cumulative probability distributions of PV/CB1 ratios in different layers. Kolmogorov-Smirnov test confirmed that the distribution of PV/CB1 ratio in L5b is different in comparison to other layers (K-S test: L2/3 vs. L5a: p= 0.73; L2/3 vs. L5b: p<0.001; L5a vs. L5b: p<0.001).

**Figure 3.**
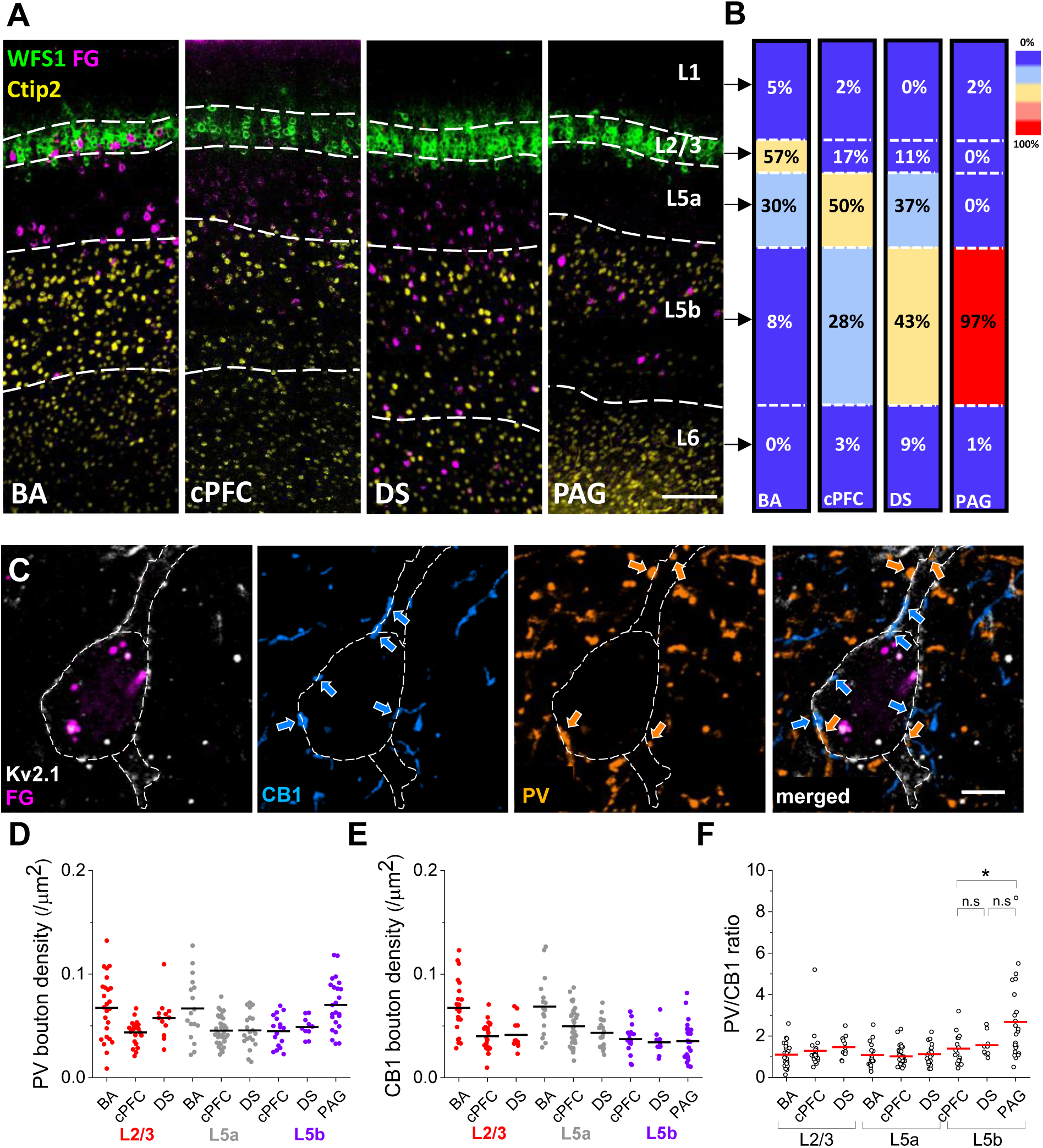
Inhibitory inputs on the perisomatic region of pyramidal cells projecting to different brain regions. (A) Multicolour confocal images taken from the mPFC showing the distribution of retrogradely labelled cells (magenta) in distinct layers. Scale bar: 100 µm. (B) Layer-specific distribution of the somata of pyramidal neurons targeting distinct brain regions. (The number of the quantified retrogradely labelled cells: Basal amygdala (BA): n=240 in 6 slices from 2 mice, Contralateral PFC (cPFC): n=1249 in 9 slices from 3 mice, Dorsal striatum (DS): n=2158 in 9 slices from 3 mice, Periaqueductal grey (PAG): n=70 in 6 slices from 2 mice). (C) Multicolour maximum z-intensity projection confocal images taken from a section immunostained for Kv2.1, Fluorogold (FG), CB1 and PV to visualise the perisomatic CB1 and PV inputs on FG-labelled neurons. Scale bar: 5 µm. (D) PV bouton density on the perisomatic surface of pyramidal neurons projecting to distinct brain regions. Red dots indicate cells located in L2/3, grey dots represent pyramidal cells located in L5a, while purple dots show pyramidal cells located in L5b. Each dot represents the PV bouton density on single pyramidal neurons. Black lines indicate means. (E) CB1 bouton density on the perisomatic membrane surface of pyramidal cells projecting to distinct brain regions. Colours, lines, dots, and numbers are the same as in (D). (F) Perisomatic PV/CB1 ratio on pyramidal cells projecting to different brain areas and located in distinct layers. Red lines indicate means. (The number of examined cells by layers and by projection brain areas: L2/3: BA: n=25, cPFC: n=22, DS: n=12; L5a: BA: n=17, cPFC: n=36, DS: n=20; L5b: cPFC: n=17, DS: n=10, PAG: n=24. BA-projecting cells were counted in 6 slices from 2 mice, cPFC-projecting cells were counted in 10 slices from 4 mice, DS-projecting cells were counted in 6 slices from 2 mice and PAG-projecting cells were counted in 4 slices from 2 mice. K-W ANOVA in L2/3 p=0.104, in L5a p=0.633 and in L5b p=0.026. In L5b M-W test showed difference between cPFC- and PAG-projecting neurons: p=0.012; DS-PAG: p=0.121, cPFC-DS: p=0.303).

In summary, our results showed that i) pyramidal cells in layer 2/3, 5a and 5b receive the vast majority of their perisomatic GABAergic inputs from two sources, and ii) the ratio of PV+/CB1+ GABAergic inputs in deep layers were higher on pyramidal cells projecting to the PAG, indicating that the primary source of perisomatic inhibition for PT neurons is PV+ cells.

### Validation of reporter protein expression in two transgenic mouse lines

Previous studies in other cortical regions have uncovered that CB1+ and PV+ axonal varicosities innervating the perisomatic region of pyramidal cells originate from two distinct kinds of basket cells (Hajos et al., 2000; Bodor et al., 2005; Hefft and Jonas, 2005; Glickfeld and Scanziani, 2006; Freund and Katona, 2007; Takacs et al., 2015; Vereczki et al., 2016). To selectively target and study the morphological characteristics of these interneurons, transgenic mouse strains can successfully be used. In our earlier studies, we used two transgenic mouse lines – BAC-CCK-DsRed (Mate et al., 2013) and BAC-PV-eGFP (Meyer et al., 2002) – to target CCKBCs and PVBCs in the hippocampus and basal amygdala (Gulyas et al., 2010; Szabo et al., 2014; Vereczki et al., 2016). Before examining the features of interneurons that can be targeted in acute slices containing the prefrontal cortical region, we validated the reporter protein expression in the mPFC in these two transgenic mouse lines. Using immunostaining, we observed that both in layer 2/3 and layer 5 the majority of strongly DsRed+ cells were immunoreactive for proCCK (55.7%, n=102 proCCK+ out of 183 DsRed+ cells), while neurons expressing weaker DsRed signals lacked detectable immunolabelling for this neuropeptide (Figure 4A-B). As this proCCK antibody stains GABAergic interneurons in the hippocampus and basolateral amygdala (Kotzadimitriou et al., 2018; Rhomberg et al., 2018), we hypothesized that neurons displaying strong DsRed signals would also be interneurons in this cortical region, while neurons expressing weaker DsRed signals might belong to a subset of pyramidal cells. This assumption has been confirmed by subsequent *in vitro* whole-cell recordings, showing that strongly DsRed+ cells were indeed interneurons (n=42 CCKBC out of 44 biocytin filled neurons). In BAC-PV-eGFP mice, our analysis demonstrated a robust co-expression of PV and eGFP (86.4%, n=686 PV+ out of 794 eGFP+ cells)(Figure 4D-E).

**Figure 4.**
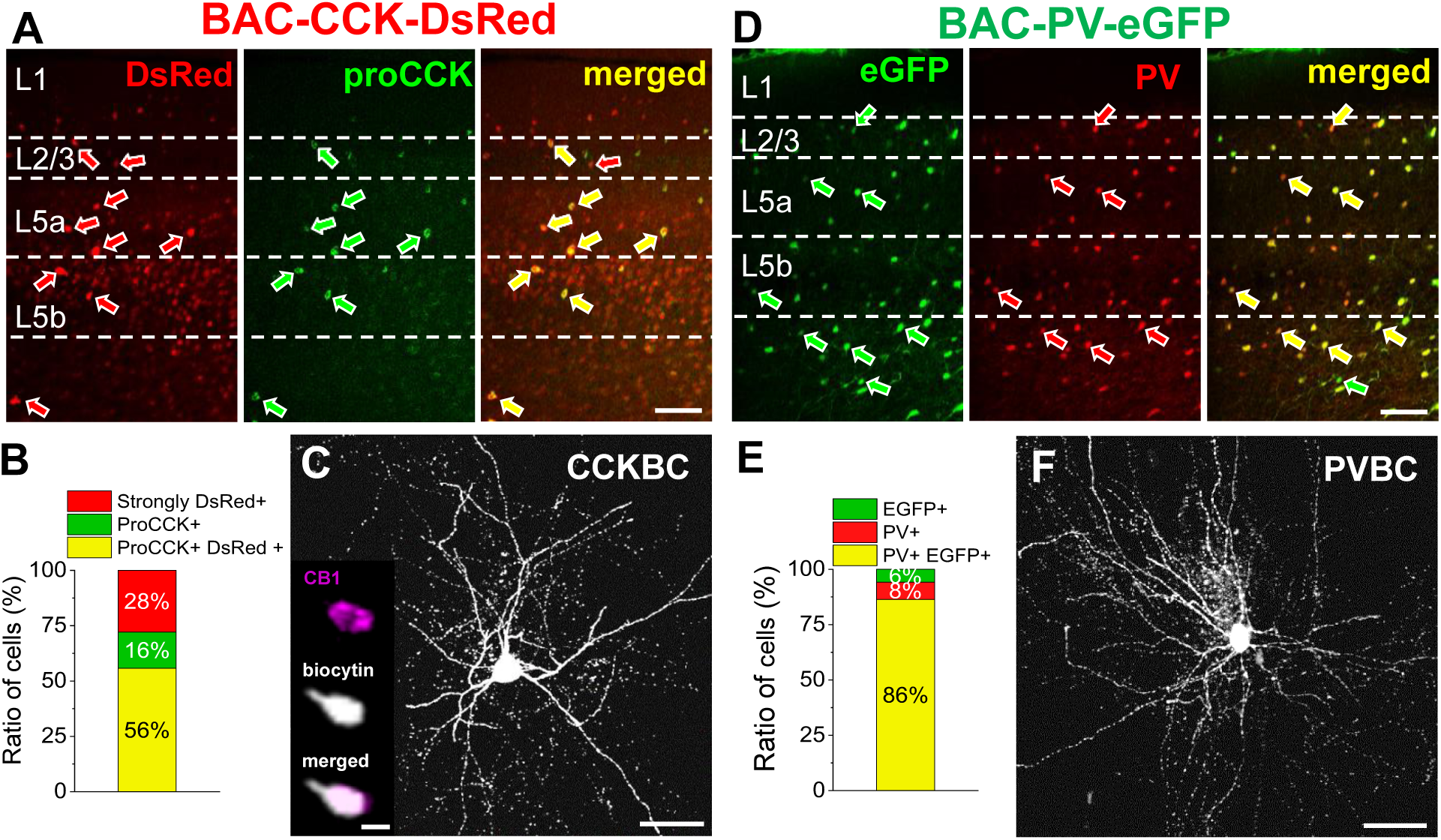
Validation of reporter protein expression in two transgenic mouse lines. (A) Multicolour confocal images taken from a mPFC-containing section prepared from a BAC-CCK-DsRed mouse. In the section proCCK was revealed by immunostaining. Red arrows show the genetically labelled DsRed+ cells, green arrows represent the proCCK-immunopositive cells, while yellow arrows indicate the co-localization of the two markers. Dashed white lines represent the boundaries of layers. Scale bar: 100 µm. (B) The ratio of counted DsRed/CCK cells within the mPFC (n=183 in 3 slices from 2 mice). (C) Maximum z-intensity projection image taken of an *in vitro* biocytin filled CCKBC. Scale bar: 50 µm. Insets present the CB1-content of a biocytin-containing bouton of the same CCKBC. Scale bar: 1 µm. (D) Multicolour confocal images taken from the mPFC of a BAC-PV-EGFP mouse. The section was immunostained for PV. Green arrows show the genetically labelled EGFP cells, red arrows represent the PV-immunolabelled neurons, while yellow arrows indicate the co-localization of the two markers. Dashed white lines represent the boundaries of layers. Scale bar: 100 µm. (E) The ratio of counted EGFP/PV cells within the PrL (n=794 in 5 slices from 2 mice). (F) Maximum z-intensity projection image taken of an *in vitro* biocytin filled PVBC. Scale bar: 50 µm.

As we have revealed earlier, CB1 immunoreactivity could be detected in axon terminals of CCKBCs sampled in the hippocampus or basal amygdala of the BAC-CCK-DsRed mouse line (Szabo et al., 2014; Andrasi et al., 2017; Veres et al., 2017). To test whether the strong DsRed+ neurons express CB1 on their axon terminals in the mPFC as well, we tested the CB1 immunoreactivity of biocytin filled neurons (Figure 4C). We found indeed, that biocytin labelled interneurons expressed this type of cannabinoid receptor on their axon terminals (n=20 CB1+ out of 20 biocytin filled neurons), i.e., strong DsRed+ neurons in the mPFC are likely CB1/CCKBCs. As PV is present both in basket and chandelier cells in cortical structures, we separated these interneurons based on the presence of cartridges formed by axonal boutons, which is a characteristic feature of chandelier cells (Somogyi, 1977). In this study, we distinguished 53 basket cells (Figure 4F) and 10 chandelier cells. Importantly, chandelier cells expressing eGFP were sampled only in layer 2/3, but not in the deeper layers, and were excluded from this study. Thus, in slices prepared from these transgenic mouse lines, basket cells expressing CCK and CB1 or PV can be readily sampled in the mPFC.

### Morphological characterization of basket cells in the mPFC

As the axonal boutons expressing PV+ and CB1+ are not evenly distributed in mPFC layers (Figure 1G, H), we tested if the dendritic and axonal tree arborization of basket cells had any layer preference. Therefore, in the next set of experiments, we aimed to investigate the morphological characteristics of basket cells in the mPFC. As biocytin was added in the pipette solution for whole-cell recordings obtained in acute brain slices, we could reconstruct the dendritic and axonal arborizations of interneurons after visualising the tracer molecules using immunostaining. Initial fluorescent microscopic examination suggested that morphologically distinct subpopulations of CCKBCs and PVBCs most probably exist in the mPFC. Therefore, we performed principal component and cluster analysis on the morphological properties of dendrites and axons to uncover subgroups within the two types of basket cells (Figure 5A_1_-A_2_). Based on the dendritic localization, CCKBCs were divided into ‘superficial’ and ‘deep’ morphological subgroups: basket cells in the former group had dendrites predominantly in layer 1, whereas basket cells in the latter one arborized primarily in layer 5a (Figure 5A_1_ and B_1_-D_2_). Regardless of the location of the CCKBCs’ somata and dendrites (Figure 5E), their axonal arbours were similar in size and were mainly restricted to layer 5a (Figure 5F). Thus, based on the location of the dendritic tree, CCKBCs could be divided into two morphological categories, however, both groups innervate pyramidal cells in the same layer of the mPFC.

**Figure 5.**
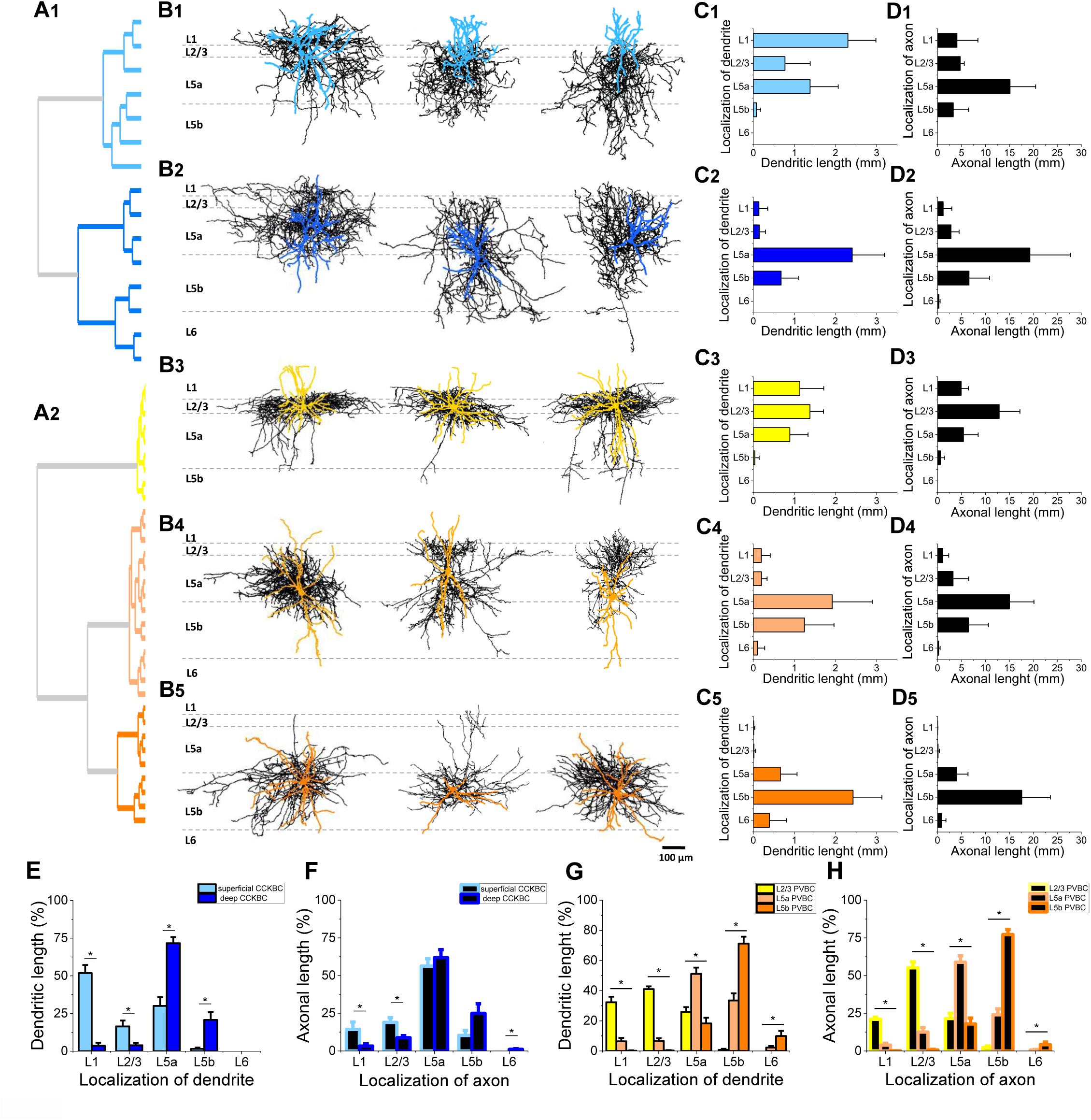
Distribution of dendritic and axonal arbour of different basket cell types within the layers of the mPFC. (A1-A2) Cluster analysis of CCKBCs determined the superficial and deep categories based on the location of their dendrites and analysis of PVBCs divided them into three subgroups based on the location of their dendrites and axons. (B1-B5) Neurolucida reconstructions of dendritic and axonal arbour of different basket cells labelled in slice preparations. Black lines represent the axons, while coloured lines show the dendritic trees, respectively. In light blue the dendrites of ‘superficial’ CCKBCs are presented (n=7 from 4 mice), while the dendrites of ‘deep’ CCKBCs are shown in dark blue (n=8 from 6 mice). With yellow the dendrites of L2/3 PVBCs (n=9 from 9 mice), with light orange the dendrites of L5a PVBCs (n=14 from 10 mice), while with bright orange the dendrites of L5b PVBCs (n=9 from 7 mice) are shown. Dashed dark grey lines represent the boundaries of the layers. Scale bar: 100 µm. (C1-C5) The distribution of dendritic lengths in the different layers. (D1-D5) The distribution of axonal lengths in the different layers. (E) Comparison of the distribution of the dendritic lengths in different layers for the two CCKBC subgroups. (M-W tests by layers: L1: p=0.001, L2/3: p=0.013, L5a: p=0.001, L5b: p=0.016) (F) Comparison of the distribution of the axonal lengths of the two CCKBC subgroups in different layers. (M-W tests by layers: L1: p=0.013, L2/3: p=0.003, L6: p=0.019) (G) Comparison of the distribution of dendritic lengths of the three PVBCs subgroups in different layers. (K-W ANOVA by layers: L1: p<0.001, L2/3: p<0.001, L5a: p<0.001, L5b: p<0.001, L6: p=0.007. *Post hoc* Dunn’s test by layers: L1: L2/3 PVBC vs L5a PVBC: p=0.002, L2/3 PVBC vs L5b PVBC: p<0.001, L5a PVBC vs L5b PVBC: n.s.; L2/3: L2/3 PVBC vs L5a PVBC: p=0.005, L2/3 PVBC vs L5b PVBC: p<0.001, L5a PVBC vs L5b PVBC: n.s.; L5a: L2/3 PVBC vs L5a PVBC: p=0.006, L2/3 PVBC vs L5b PVBC: n.s., L5a PVBC vs L5b PVBC: p<0.001; L5b: L2/3 PVBC vs L5a PVBC: p=0.03, L2/3 PVBC vs L5b PVBC: p<0.001, L5a PVBC vs L5b PVBC: p=0.008; L6: L2/3 PVBC vs L5a PVBC: n.s., L2/3 PVBC vs L5b PVBC: p=0.005, L5a PVBC vs L5b PVBC: n.s.) (H) Comparison of the distribution of axons of the three PVBCs subgroups in different layers. (K-W ANOVA by layers: L1: p<0.001, L2/3: p<0.001, L5a: p<0.001, L5b: p<0.001, L6: p=0.005. *Post hoc* Dunn’s test by layers: L1: L2/3 PVBC vs L5a PVBC: p=0.005, L2/3 PVBC vs L5b PVBC: p<0.001, L5a PVBC vs L5b PVBC: n.s.; L2/3: L2/3 PVBC vs L5a PVBC: p=0.007, L2/3 PVBC vs L5b PVBC: p<0.001, L5a PVBC vs L5b PVBC: p=0.04; L5a: L2/3 PVBC vs L5a PVBC: p<0.001, L2/3 PVBC vs L5b PVBC: n.s., L5a PVBC vs L5b PVBC: p<0.001; L5b: L2/3 PVBC vs L5a PVBC: p=0.02, L2/3 PVBC vs L5b PVBC: p<0.001, L5a PVBC vs L5b PVBC: p=0.01; L6: L2/3 PVBC vs L5a PVBC: n.s., L2/3 PVBC vs L5b PVBC: p=0.004, L5a PVBC vs L5b PVBC: n.s.)

In comparison to CCKBCs, PVBCs showed a larger diversity. Based on the location of their dendrites and axons, cluster analysis separated PVBCs into three subgroups (Figure 5A_2_ and B_3_-D_5_). The layer 2/3 PVBCs spread approximately half of their axons within the layer 2/3 (Figure 5H), while their dendrites were located in layer 1, 2/3 and 5a with similar ratios (Figure 5C3, G). The layer 5a PVBCs and layer 5b PVBCs extended the majority of their dendrites and axons in layer 5a and layer 5b, respectively, with the vast majority of processes in the same layer where their soma was located (Figure 5G-H). These observations indicate that PVBCs mostly innervate neurons in the same layers where they receive most of their inputs.

Next, we compared the characteristics of the dendritic and axonal arbours between the two basket cell types. First, when comparing the total length of the dendrites and axons we found no difference between CCKBCs and PVBCs (Figure 6A-B), yet PVBCs spread their dendrites further from the soma than CCKBCs (Figure 6H_1_). Although both cell types showed similar axonal length, the axon of PVBCs had significantly more axonal nodes resulting in a lower length/nodes ratio in comparison to CCKBCs (Figure 6C-D), indicating a more branchy axonal cloud. Second, for both basket cell types we correlated the total axonal length with the total dendritic length: a significant relationship was observed between these two parameters for PVBCs, but not for CCKBCs (Figure 6E_1-_E_2_). These results imply that those PVBCs that are collecting more inputs within the microcircuit may inhibit more pyramidal cells. Third, we examined the fine structure of their dendrites and axons. PVBCs emitted more primary, secondary and tertiary dendrites, while CCKBCs had longer higher-order dendritic segments (Figure 6F). Regarding the axons, we observed using Sholl analysis (Figure 6G_1_-G_2_) that the reconstructed CCKBCs had longer axons 250-400 µm from their somata than PVBCs, implying that the output structure of the two basket cells is distinct (Figure 6H_2_). Based on detailed morphological analysis, these findings show that in comparison to CCKBCs, PVBCs have more complex dendritic and axonal trees, which arborize mainly within 200 µm from the somata (Figure 6H_1_-H_2_). These structural features may be in line with their different roles played in mPFC operation.

**Figure 6.**
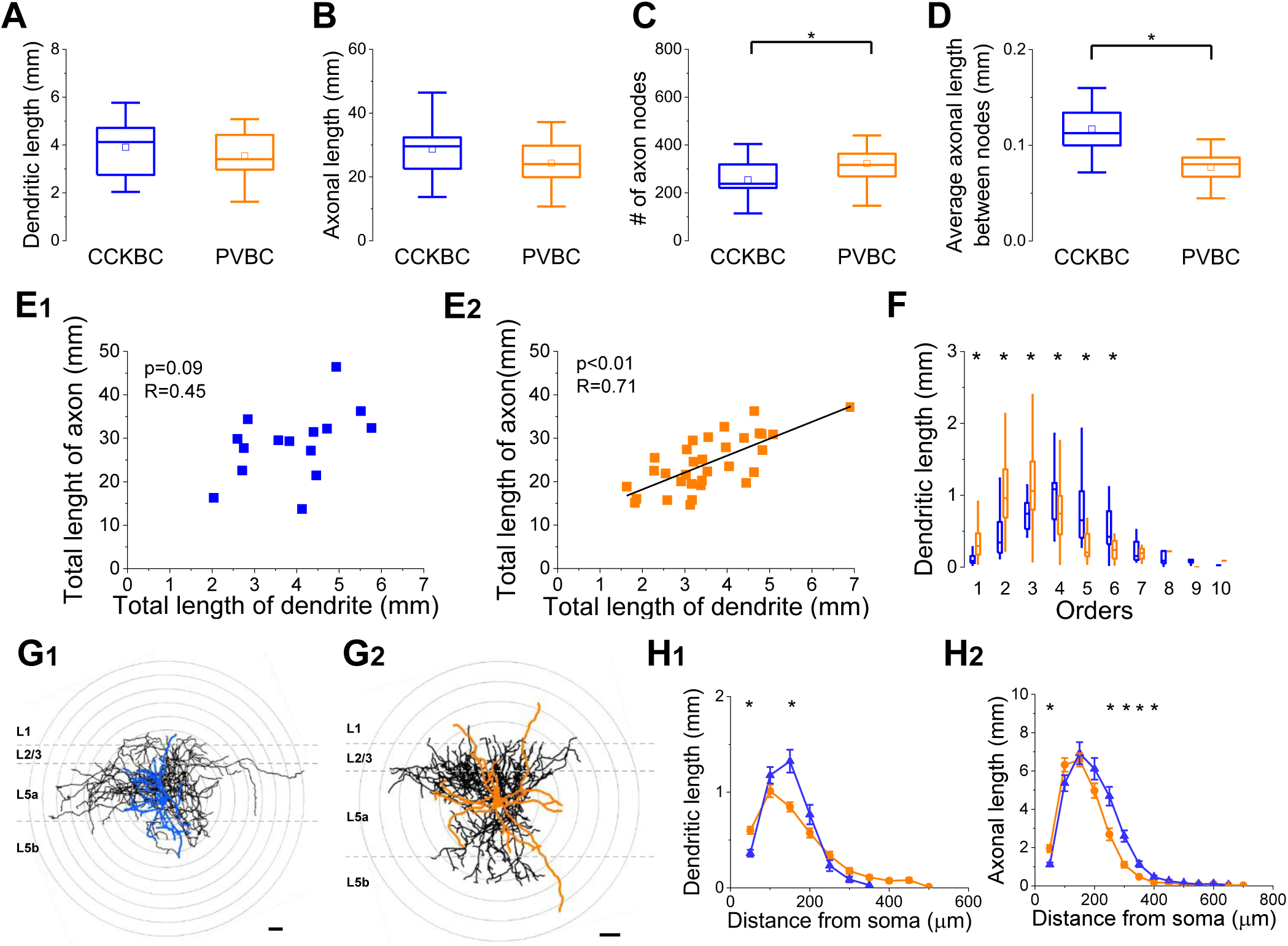
Comparison of the dendritic and axonal arborisations of CCKBCs and PVBCs. (A-D) Box chart comparison of the total dendritic and axonal length, total number of axonal nodes and average axonal length between nodes. The mean (small open square), the median (continuous line within the box), the interquartile range (box) and the 5-95% values (ends of whiskers bar) are plotted. (Statistical comparisons were performed with M-W test: (C) p=0.022; (D) p<0.001.) (E_1_-E_2_) Total axonal length as a function of the total dendritic length. Each dot corresponds to the values of a reconstructed BC. A significant relationship was found between these two values in case of PVBCs (Pearson’s r: R=0.71; p<0.001). (F) Comparison of the dendritic length as a function of the dendritic order. (M-W test: 1. order: p<0.001; 2. order: p<0.001; 3. order: p=0.019; 4. order: p=0.023; 5. order: p<0.001; 6. order: p=0.018) (G_1_-G_2_) Neurolucida reconstructions of dendritic and axonal tree of two example BCs labelled in slice preparations. Black lines represent the axons, while coloured lines show the dendritic trees. Concentric grey circles drawn on the reconstructions illustrate the 50 µm radii used for Sholl analysis. Dashed grey lines represent the boundaries of the layers in the mPFC. Scale bar: 50 µm. (H_1_) The dendritic length as a function of the distance from the soma. (M-W test showed differences at 0-50 µm distance from the soma (p<0.001) and at 100-150 µm (p=0.001)) (H_2_) The axonal length as a function of the distance from the soma. (M-W test showed differences at 0-50 µm distance from the soma (p=0.002), at 200-250 µm (p<0.001), at 250-300 µm (p<0.001), at 300-350 µm (p<0.001) and at 350-400 µm (p=0.002))

### Postsynaptic target distribution of basket cells within the mPFC

To verify that interneurons sampled in the two transgenic mouse lines indeed provide the perisomatic innervation of neurons in the mPFC as in other cortical circuits (Glickfeld and Scanziani, 2006; Foldy et al., 2007; Kohus et al., 2016; Veres et al., 2017), we investigated their postsynaptic target distribution. Therefore, we examined the ratio of axonal varicosities forming close appositions with the Kv2.1-labelled perisomatic region to those avoiding them (Figure 7A-B). The analysis showed that around 55% of the boutons of both CCK+ and PV+ interneurons targeted the somata and proximal dendrites (Figure 7D), an observation which proves that our recorded neurons are basket cells indeed. Importantly, although the ratio of axonal varicosities contacting the somata was fairly similar between the two basket cell types and between the morphologically distinct subpopulations (Figure 7C-E), there was a substantial variance in the ratio of boutons innervating the proximal dendritic segments within both main groups (Figure 7E). In addition, we addressed the question whether there is a difference in the number of perisomatic contacts originating from CCKBCs and PVBCs at the single cell level. Therefore, we examined the bouton numbers on approximately 20 Kv2.1-labelled neurons located within 150-200 µm from the in vitro filled basket cells. Interestingly, we observed that single Kv2.1-labelled neurons received more boutons from a PVBC than from a CCKBC (Figure 7F). Overall, our findings show that i) CCK+ and PV+ interneurons (that differ from chandelier cells) preferentially target the perisomatic region of mPFC neurons, i.e., they are indeed basket cells and ii) there are no interneurons expressing CCK or PV in the mPFC that innervate preferentially dendrites like in the hippocampus (Halasy et al., 1996; Cope et al., 2002; Somogyi and Klausberger, 2005; Szabo et al., 2014).

**Figure 7.**
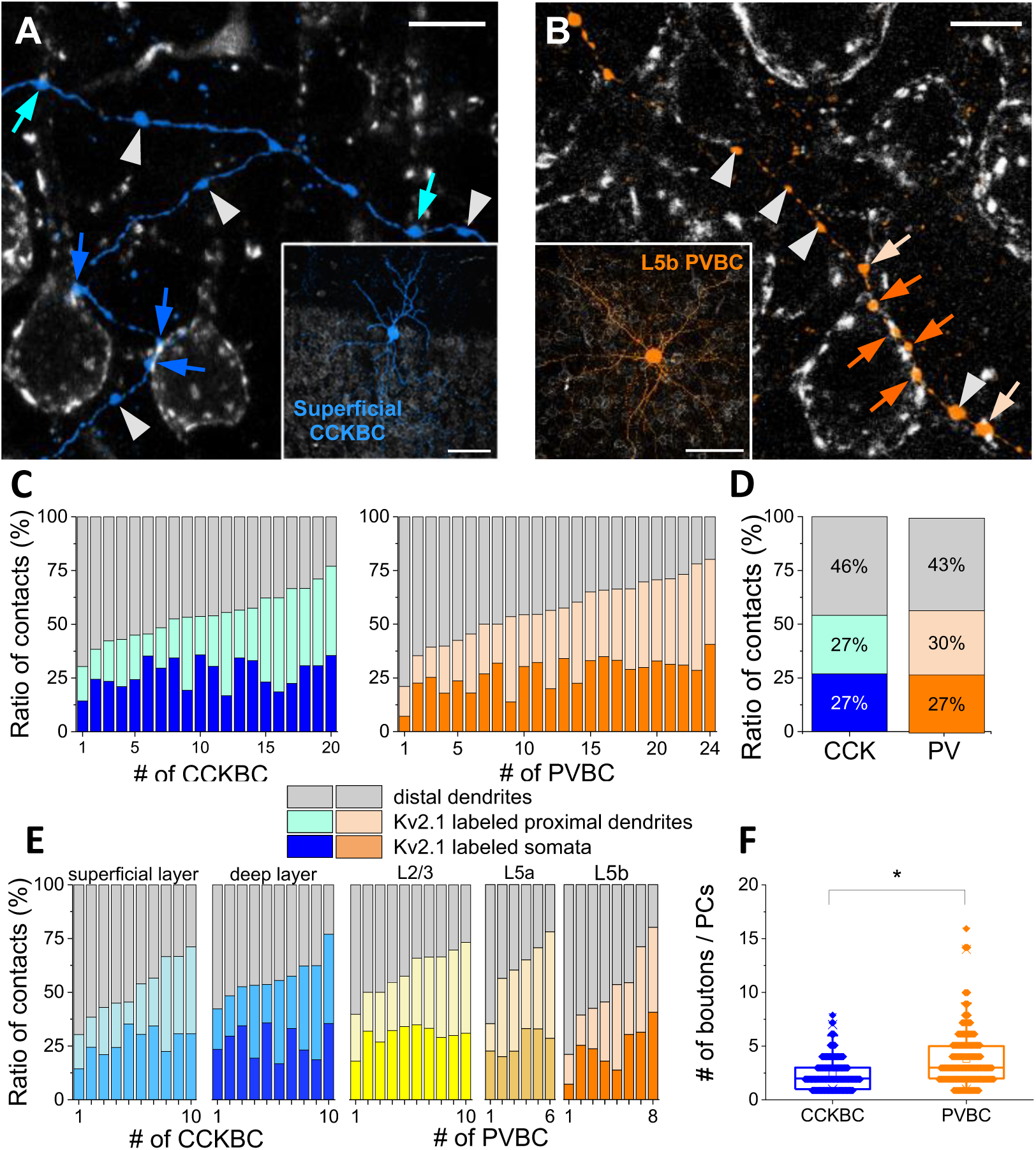
Postsynaptic target distribution of basket cells within the mPFC. (A) High magnification multicolour maximum z-intensity projection confocal image shows biocytin-filled boutons around Kv2.1-labelled neurons, dark blue arrows point to boutons forming contacts on the somata, light blue arrows indicate the proximal dendrite-targeting boutons, while grey arrows show varicosities that presumably target distal dendrites (i.e., avoid Kv2.1 labelled profiles). Scale bar: 10 µm. Inset shows the *in vitro* biocytin-filled ‘superficial’ CCKBC, whose axons are shown at the higher magnification. Scale bar: 100 µm. (B) High magnification multicolour maximum z-intensity projection confocal image shows biocytin-filled boutons around Kv2.1-labelled neurons, bright orange arrows point to boutons forming close contacts on a soma, light orange arrows indicate the proximal dendrite-targeting boutons, while grey arrows show varicosities that likely target distal dendrites (i.e., avoid Kv2.1 labelled profiles). Scale bar: 10 µm. Inset shows the *in vitro* biocytin-filled L5b PVBC, whose axon collaterals are shown at the higher magnification. Scale bar: 100 µm. (C) Ratio of the boutons of CCKBCs and PVBCs forming close contacts on Kv2.1-immunostained perisomatic regions of neurons. Each column represents a single BC (n=20 CCKBC and n=24 PVBC were examined. M-W test showed no difference in the ratio of boutons on Kv2.1-immunolabelled soma between CCKBC and PVBC (p=0.99)). (D) Average ratio of biocytin-filled boutons forming contacts on Kv2.1-immunolabelled profiles or on unlabelled targets (pooled data from C). (E) Ratio of the boutons of the basket cell subgroups. Each column represents a single basket cell (n=3560 boutons of 10 ‘superficial’, n=3518 boutons of 10 ‘deep’ CCKBC were examined. M-W test showed no difference in the ratio of boutons on Kv2.1-immunostained somas between distinct groups of CCKBCs (p=0.97). n=2838 boutons of 10 L2/3, n=1986 boutons of 6 L5a and n=1995 boutons of 8 L5b PVBC were examined. K-W ANOVA showed no difference in the ratio of boutons on Kv2.1-immunostained somas between distinct groups of PVBCs (p=0.217)). (F) Comparison of the number of the perisomatic contacts received by single Kv2.1-labelled neurons from an individual CCKBC and PVBC. The mean (small open square, CCKBC: 2.46; PVBC: 3.82), the median (continuous line within the box, CCKBC: 2; PVBC: 3), the interquartile range (box, CCKBC: 2; PVBC: 3) and the 5/95% values (ends of whiskers bar) are plotted. 211 Kv2.1-labelled neurons innervated by 520 boutons from 10 CCKBC and 199 Kv2.1-labelled neurons innervated by 761 boutons from 10 PVBC were examined (M-W test: p<0.001).

### PVBCs are preferentially targeted by thalamic and amygdalar inputs in layer 5a and 5b, respectively

We have shown that PVBCs preferentially innervate the same layer in the mPFC where their somata are located (Figure 5). As different mPFC afferents typically target some layers, but not others (Anastasiades and Carter, 2021), this raises the possibility that PVBCs are capable of mediating feedforward inhibition in a layer-specific manner if they are preferentially targeted by extra-mPFC inputs in a given layer. To test this hypothesis, we investigated the innervation of PV+ interneurons by afferents originating either from the BA, midline thalamus (Thal) or lateral entorhinal cortex (LEnt). We combined anterograde trans-synaptic viral labelling with immunostaining and determined the ratio of PV+ interneurons that did and did not receive monosynaptic inputs from distinct projections in each layer. To obtain anterograde trans-synaptic labelling two approaches were used. AAV1-hSyn-Cre was injected into the Ai6 reporter line resulting in ZsGreen1 expression in neurons that receive monosynaptic innervation (Figure 8A). In these experiments, PV+ interneurons were revealed by immunostaining. The other approach involved PV-Cre mice, in which AAV1-EF1a-DIO-ChETA-eYFP was injected, directly revealing monosynaptically innervated PV+ interneurons. By using these approaches, we were able to visualise those neurons in the mPFC that receive monosynaptic innervation from the injected area (Zingg et al., 2017) (Figure 8B). Since both strategies gave rise to similar ratio of trans-synaptically labelled PV+ cells, this confirms that the labelling success was not dependent on the AAV1 or mouse strains used in these investigations (Figure 8C). Therefore, we pooled the data from the two approaches. Next, we compared the distribution of PV+ interneurons in the different layers that received inputs from the BA, Thal or LEnt input in the distinct layers with the distribution of all PV-immunolabelled interneurons in the mPFC (Figure 8D). We observed that afferents from the BA and Thal innervated PV+ interneurons in a layer specific manner, but not the input arriving from LEnt. In case of BA inputs, a higher ratio of PV+ interneurons was labelled with AAVs in layer 5b compared to the relative distribution of all PV-immunostained interneurons, and the same was true in case of Thal inputs in layer 5a (Figure 8E). As we found no PV+ chandelier cells in layer 5a and 5b of the mPFC in line with a recent study (Wang et al., 2019), these data imply that excitation from the amygdala and thalamus can drive PVBCs-mediated feedforward inhibition in the mPFC in a layer specific manner.

**Figure 8.**
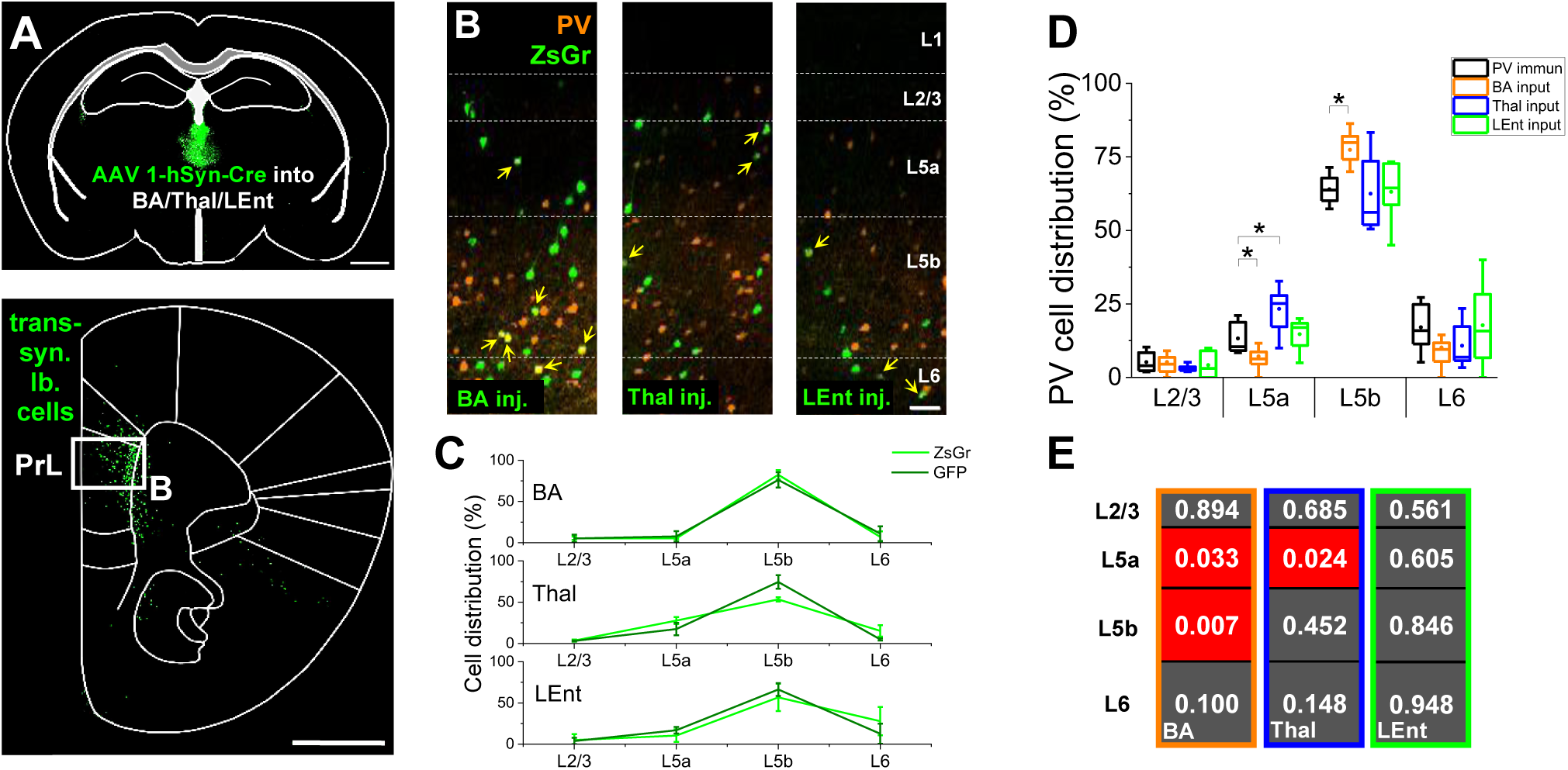
PV-expressing interneurons in the mPFC are preferentially innervated by thalamic (Thal) and amygdalar (BA) afferents in layer 5a and 5b, respectively. (A) Upper panel shows the schematic representation of AAV1-hSyn-Cre virus injection into Thal. Bottom panel represents the localization of trans-synaptically labelled cells within the mPFC. Scale bar: 1 mm. (B) Multicolour maximum z-intensity projection confocal images taken from the prelimbic (PrL) area of Ai6 mice injected with AAVs into the BA, Thal and lateral entorhinal cortex (LEnt). Immunostaining was used to visualise PV interneurons. Yellow arrows point to ZsGreen1 (ZsGr) and PV double positive cells. Scale bar: 50 µm. (C) Comparison of PV-immunolabelled ZsGr- or GFP-positive cell distribution in each layer of Ai6 and PV-Cre mouse lines upon AAV injections into different areas. Chi-Square Homogenity test showed no differences between mouse lines. (BA: χ2=0.09, p=0.99; Thal: χ2=1.86, p=0.6; LEnt: χ2=0.2, p=0.98) (D) Box chart comparison of PV cell ratios in different layers. PV-immunolabelled interneurons are shown in black (n=1283 PV immunopositive cells in 12 slices from 7 mice), the BA input receiving PV cells in orange (n=570 cells in 45 slices from 10 mice), Thal input receiving PV cells in blue (n=731 cells in 38 slices from 7 mice), while LEnt input receiving PV cells in green (n=187 cells in 29 slices from 6 mice). The mean (small open square), the median (continuous line within the box), the interquartile range (box) and the 5/95% values (ends of whiskers bar) are plotted. (E) p values of M-W test comparison between the ratios PV cells receiving a given extra-mPFC inputs versus PV-immunolabelled cells in each layer (Black versus coloured bars in panel (D)). Red background indicates significant differences.

## Discussion

In this study, GABAergic inputs innervating the perisomatic region of neurons in the PrL subregion of the mPFC were examined. We identified the sources of these inputs and compared the morphological features of interneurons that provide the vast majority of perisomatic inhibitory innervation on the soma and proximal dendrites. Besides the layer definition in the mPFC our main findings are the followings: (1) the perisomatic region of pyramidal cells in the mPFC is innervated mostly by PV+ or CB1+ GABAergic inputs. (2) These inhibitory inputs originate from PVBCs and CCKBCs. (3) At the population level, more than 50% of axonal boutons of both basket cell types contacted the perisomatic region. However, at the single cell level, PVBCs innervated the perisomatic region via more boutons than CCKBCs. (4) Most importantly, we found that PVBCs are innervated by afferents originating from the BA and Thal in a layer-specific manner.

Although several studies have examined the cytoarchitectural and connectivity properties of the mPFC, the layers of this brain region are still ill-defined and show inconsistency in the publications (Clarkson et al., 2017; Lu et al., 2017; Wang et al., 2019; Anastasiades and Carter, 2021). Recent studies have established probably the most precise method to define a cortical layer by using single-cell transcriptomics (Tasic et al., 2018; Wang et al., 2018; Ortiz et al., 2020), unfortunately it is hard to adopt this methodology to acute brain slices used in electrophysiological recordings. Additionally, as the thickness of layers in the mPFC changes continuously in dorso-ventral and anterior-posterior directions, the measurements of the distance from the pia are not the most reliable way to define the layers. Therefore, we introduced an approach and defined the layers of the mPFC by immunostaining using a combination of antibodies developed against markers that have been used in other cortical areas as well (Cruikshank et al., 2001; Arlotta et al., 2005; Luuk et al., 2008; Hisaoka et al., 2010).

Similarly to other cortical studies, we observed that the perisomatic region of neurons is innervated mostly by GABAergic inputs from two sources (Freund and Katona, 2007; Takacs et al., 2015; Vereczki et al., 2016). Our findings revealed that around 90% of inhibitory inputs on the perisomatic membrane surface is originating from PV+ or CB1+ axonal varicosities. Interestingly, there was a layer specific difference in the ratio of perisomatic PV+/CB1+ inputs on randomly sampled neurons (Figure 2D) and when examined on defined pyramidal cell populations projecting to specific areas (Figure 3F). Thus, additionally to the observations that the two types of basket cells contribute mainly to perisomatic inhibition in the mPFC, it seems that pyramidal cells located in deeper layers are innervated by more inputs from PVBCs. A previous study using a combination of optogenetics with slice physiology proposed that PVBCs preferentially innervate PT neurons in comparison to intratelencephalic (IT) neurons (Lee et al., 2014) which is in line with our anatomical findings. However, these data contradict those obtained with paired recordings in the frontal cortex, suggesting that IT and PT neurons receive similar strength of inhibition from putative PV interneurons (Morishima et al., 2017). PT neurons in the mPFC are also innervated by CCKBCs, as we found in the present study. This finding is in contract to our observations obtained in the somatosensory cortex, where we detected only negligible perisomatic inputs from CCKBCs onto the PT neurons (Bodor et al., 2005). Altogether, these results may support the hypothesis that in primary sensory cortices PT neurons are controlled solely by PVBCs, whereas in higher order cortical areas, the pyramidal cells are under the regulation of both basket cell types (Toyoda, 2020), an idea that needs to be specifically tested.

Several studies have examined the features of these two basket cell types separately in rodent neocortex (Kawaguchi and Kubota, 1998; Lagler et al., 2016; Miyamae et al., 2017), yet the comparison of their morphological characteristics has not been conducted in the mouse mPFC. Our morphological analysis and comparison revealed structural differences between the two basket cell types. Although the total dendritic and axonal length of the two basket cell types were similar, we found differences in the structure of their dendritic and axonal arborizations. Interestingly, similar differences were observed in the basal amygdala too, suggesting that basket cells might not possess brain region specific structural features (Vereczki et al., 2016), but have a more uniform morphological structure. We have also found that distinct basket cell types could be divided into further subgroups based on the location of their somata, dendrites and axons. Such morphological subgroups of CCKBCs have not been proposed so far, yet distinct morphological types of PVBCs were described in previous studies (Jiang et al., 2015; Lagler et al., 2016; Miyamae et al., 2017) without any systematic comparison. As PVBCs play a pivotal role in the generation of gamma oscillations (Sohal et al., 2009; Gulyas et al., 2010), the layer-specific arborization of these GABAergic interneurons may ensure the independent oscillatory activities in different layers, a hypothesis which is in accord with recent in vivo findings (Senzai et al., 2019). Thus, the structural features of PVBCs may allow the neural computation to be performed synchronously across cortical layers or restricted to a given layer, depending on the primary excitatory inputs driving PVBC spiking.

Multicolour labelling allowed us to analyse the target distribution of distinct basket cells. We found that on average more than 50% of the biocytin-filled boutons of both basket cell types innervated the perisomatic region of pyramidal cells. Additionally, a large variability was observed in the ratio of contacts forming close appositions with the perisomatic membrane surface in both cases. Interestingly, this variance was due to the difference in the innervation of the proximal dendrites, as the ratio of boutons innervating the somata was similar within and between the basket cell types. These findings are opposite to those observed in the BA, where the ratio of boutons contacting the somata showed a large variance, whereas the proportion of boutons on proximal dendrites was rather similar (Veres et al., 2017). Thus, the innervation strategy of basket cells seems to be distinct in different cortical structures, which may relate to the morphological differences of cortical principal neurons (i.e., stellate like appearance of principal neurons in the BA vs. pyramidal neurons in the mPFC). Moreover, between the two extreme values, the ratio of perisomatic innervation on the soma vs the proximal dendrite varied continuously, similarly to that observed in the BA (Veres et al., 2017). Furthermore, when we counted the number of boutons innervating single pyramidal cells in the vicinity of basket cells, we found that PVBCs targeted pyramidal cells with significantly more boutons than CCKBCs, similarly to that found in the BA (Vereczki et al., 2016). However, this difference in the number of boutons may not be reflected in the efficacy of inhibition (Veres et al., 2017).

Our morphological analyses revealed that PVBCs extended mostly their dendrites and axons in the same layer where their somata was located (Figure 5A_3_-C_5_). In addition, previous data demonstrated that extra-mPFC afferents from distinct brain regions show a layer-specific distribution within the mPFC (Parnaudeau et al., 2013; Oh et al., 2014; McGarry and Carter, 2016; Anastasiades and Carter, 2021). Therefore, based on these observations, we hypothesized that PV+ interneurons (which are predominantly basket cells in the mPFC based on our study) can be innervated preferentially by inputs from the BA, Thal and LEnt cortex in a layer-specific manner. In line with this idea, we found that thalamic and amygdalar inputs favoured PVBCs in layer 5a and 5b, respectively (Figure 8D-E). Thus, our anatomical findings suggest that excitatory inputs from the BA and Thal, but not from the LEnt cortex can mediate feedforward inhibition via PVBCs in the mPFC in a layer-specific manner.

Taken together, our results demonstrated the presence of morphologically different basket cells in the mPFC microcircuits. Furthermore, PVBCs in different layers are preferentially contacted by distinct extra-mPFC afferents. Thus, PVBCs innervating pyramidal cells in their vicinity have the possibility to control mPFC function in a layer-specific manner. In contrast, CCKBCs with extended axonal arborization can provide cross-layer inhibition in the mPFC. The different innervation strategy of the two basket cell types may play a key role in controlling neural operation within the individual layers and across the layers of the mPFC.

## Author contributions

Designed experiments: JMV, NH

Performed experiments: PNP, JMV, ZsF, MRK, FW, BB, ZsR

Analysed data: PNP, JMV

Supervised the project: JMV, NH

Wrote the paper: PNP, JMV, NH

## Acknowledgements

We acknowledge financial support from the Hungarian Brain Research Program (2017-1.2.1-NKP-2017-00002) and National Research, Development and Innovation Office (K131893). The authors are grateful to Péter Földi, Éva Krizsán, Orsolya Papp and Erzsébet Gregori for their excellent technical assistance. We also thank László Barna, the Nikon Microscopy Center at the Institute of Experimental Medicine, Nikon Austria GmbH, and Auro-Science Consulting, Ltd., for kindly providing microscopy support.

